# Moment-Based Estimation of State-Switching Rates in Cell Populations

**DOI:** 10.1101/2022.01.06.475260

**Authors:** Michael Saint-Antoine, Abhyudai Singh

## Abstract

In isogenic cell populations, cells can switch back and forth between different gene expression states. These expression states can be biologically relevant. For example, a certain expression state may cause a tumor cell to be resistant to treatment, while another state may leave it vulnerable to treatment. However, estimating the rates of state-switching can be difficult, because experimentally measuring a cell’s transcriptome often involves destroying the cell, so it can only be measured once. In this paper, we propose a computational method to estimate the rate of switching between expression states, given data from a Luria-Delbrück style fluctuation test that is experimentally simple and feasible. We then benchmark this method using simulated data to test its efficacy, with varying assumptions made about cell cycle timing distribution in the simulations.

## I. INTRODUCTION

Even in clonal groups of cells that share the same genetic code, cells sometimes exhibit different phenotypic characteristics, caused by different patterns of gene expression [1]. This phenomenon is referred to as “cellular heterogeneity”, and can arise due to non-genetic factors affecting gene expression, such as epigenetic regulation and stochastic variability in the processes of transcription and translation [2]-[14]. Recent developments in experimental technology, such as single-cell RNA-seq (scRNA-seq) have made it feasible to study the transcriptomic profiles of individual cells, yielding new insights about cellular heterogeneity at the single-cell level [15]. Heterogeneity in cell populations is known to play a role in many biological contexts, including drug response in cancer [16]–[18], latency in HIV cells [19], [20], immune response in epithelial tissue [21], determination of cell fate in genetically identical populations [22]–[31], and “bet-hedging” responses to environmental stresses in bacteria [32], [33]. For more information, an excellent review of the study of cellular heterogeneity can be found in the Altschuler and Wu (2010) paper “Cellular Heterogeneity: When Do Differences Make a Difference?” [34].

We are especially interested in the phenotypic state-switching dynamics associated with transient cellular heterogeneity – that is, when cells switch back and forth between different phenotypic states. A fascinating example of this is related to drug resistance in cancer. Recent research has uncovered a phenomenon in which melanoma cells switch back and forth between a common drug-sensitive state and a rare pre-resistant state [35], [36]. When the drug is administered, cells in the sensitive state die off, while cells in the pre-resistant state undergo a process in which they become “locked in” to a state of permanent drug resistance, and no longer switch back to the sensitive state [35], [36]. Similar phenomena have been observed in many different types of cancer [37]–[40] and their significance for treatment scheduling has been explored in theoretical models [41], [42].

In cases like this, where cells switch back and forth between different phenotypic states, it can be quite challenging to estimate the rates of switching between the states. This is because experimentally measuring a cell’s transcriptome often involves destroying the cell, so it can only be measured once for each cell. In this paper, we will propose a computational method to estimate the rate of switching between expression states that is experimentally simple and feasible.

## II. Experimental Setup

In 1943, researchers Salvador Luria and Max Delbrück used a fluctuation test experiment to investigate whether bacterial resistance to T1 phage occurred spontaneously or was induced by the virus [43]. The fluctuation test experiment involved growing out many colonies of bacteria, and then exposing each colony to the phage. Luria and Delbrück counted the number of surviving bacteria in each colony, and studied their distribution. They realized that if resistance were being induced by the virus, the numbers of surviving bacteria would follow a poisson distribution. However, they found instead that the distribution of surviving bacteria showed much greater variance, suggesting that resistance to the virus was caused by a mutation that occurred prior to exposure to the virus. The results of this experiment were evidence of Darwinian selection acting on bacteria by way of random mutations that confer a fitness advantage. Since then, the experiment has been analyzed theoretically, with researchers deriving probability distributions for the number of resistant cells based on different biological assumptions [44]–[46]. An excellent review of some of the theoretical developments is available in the Zheng 1999 paper “Progress of a half century in the study of the Luria-Delbrück distribution” [47].

We can use a modified fluctuation test to study the rate at which cells switch between transient phenotypic states. Consider a population of cells that switch back and forth between two expression states, an ON state and an OFF state. This modified fluctuation test will involve a workflow similar to the original Luria-Delbrück experiment. First, several individual cells will be isolated from the population. Then, each of these isolated cells will be grown out into its own colony. After a set amount of time, the fraction of ON cells in each colony will be recorded. This may be done by transcriptome sequencing, using methods such as RNA-Seq. Or, in cases where the ON state corresponds to resistance to a drug or virus, the colonies could simply be exposed to the drug or virus and the survivors counted. Please see Figure 1 for a graphical representation of this experiment.

**Fig. 1.**
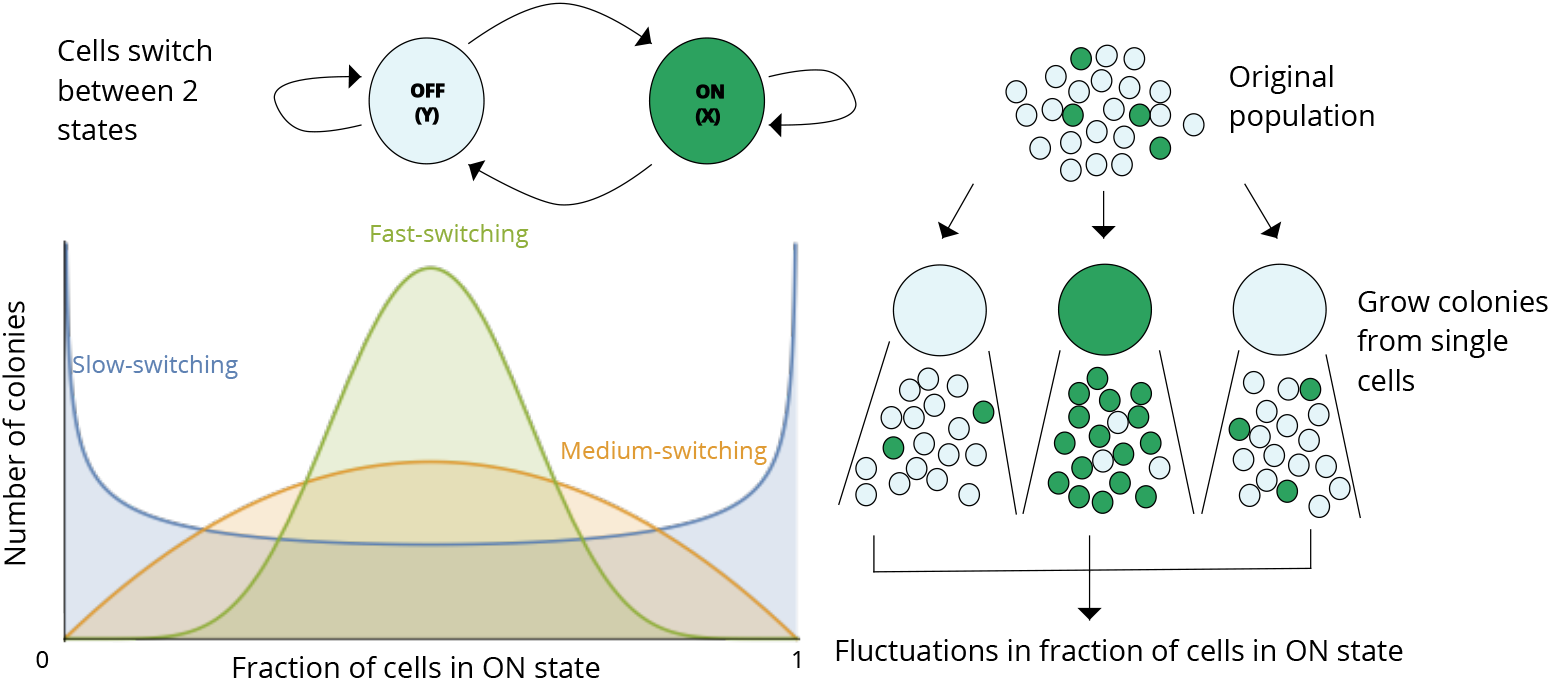
Workflow of the modified fluctuation test experiment. We begin with an original population of cells that switch back and forth between an OFF (*Y*) state and an ON (*X*) state. We isolate many individual cells, and grow each one out into a colony. After some time, we measure the fraction of cells in the ON state in each colony, and calculate the *CV*^2^ in these fractions across colonies. All else equal, populations of cells with faster state-switching rates will tend to have lower variation in these fractions than populations of cells with slower state-switching rates, since the faster-switching populations will tend to converge back to the steady state fraction of ON cells in the original population more quickly.

Once we have recorded the fraction of ON cells in each colony, we can then look at the coefficient of variation squared (*CV*^2^) of these fractions in order to infer the rates of switching between the ON and OFF states. Slow rates of switching between ON and OFF states will tend to yield higher *CV*^2^, while faster rates of switching will tend to yield lower *CV*^2^. The intuition here is that each of the colonies will eventually converge back to the original population average of cells in the ON state. However, faster-switching populations will converge back more quickly, leading to lower *CV^2^* in fractions across colonies, and the reverse is true for slower-switching populations.

Variations of this modified fluctuation test have previously been used to study HIV [19] and cancer [41], [48], [49]. The test has also been studied theoretically through mathematical analysis [50]. However, to our knowledge, there has not yet been an attempt to benchmark this method on simulated “gold standard” data, for which the true switching rates are already known. As we will see in the next section, the mathematical formula used in this technique is an inexact approximation. So, it is a pressing issue to validate this method through a benchmarking process so that we can be confident in its accuracy as it continues to be used on real-world experimental datasets.

## III. Formula Derivation

In this section, we will derive a formula for the *CV*^2^ of the fractions of ON cells across colonies in the modified fluctuation test experiment. Let us again consider a population in which cells switch back and forth between an OFF state, which we will call *Y*, and an ON state, which we will call *X*. Please see Figure 2 for a graphical representation of these dynamics.

**Fig. 2.**
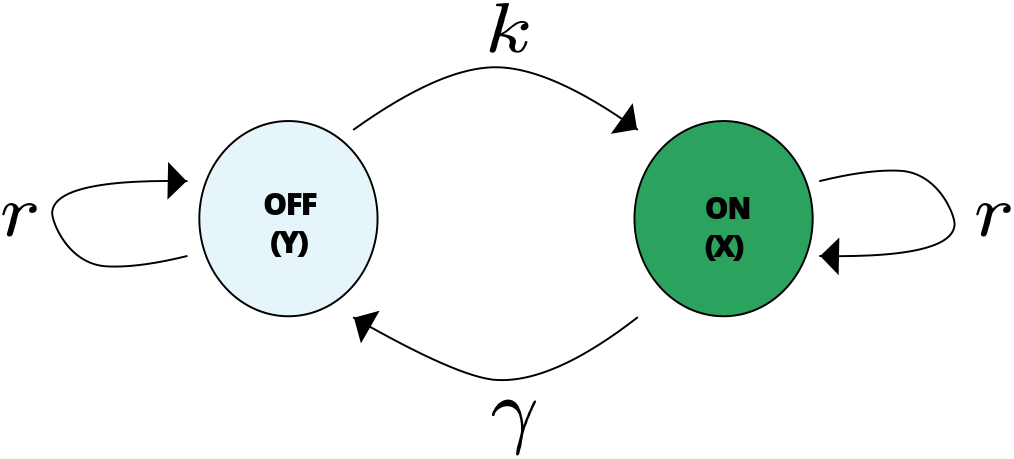
Cells switch back and forth between states OFF (*Y*) and ON (*X*). They switch from OFF to ON with rate *k*, and from ON to OFF with rate *γ*. Cells in both states proliferate with rate *r*.

We assume that the cell cycle time is an exponentially-distributed random variable with a mean generation time of 1/*r*. Hence, starting from a single cell the population grows exponentially with rate *r*. We make two simplifying assumptions: i) The proliferation rate of a cell is the same irrespective of the cellular state; ii) The population remains in the exponential growth phase during the timespan of the experiment. Cells in the OFF state transition to the ON state with rate *k*, and switch back to the OFF state with rate *γ*. So, the expected amount of time a cell will spend in the OFF state is 1/*k*, and the expected amount of time a cell will spend in the ON state is 1/*γ*. Since we have defined the cell proliferation rate as *r*, the expected number of generations a cell will spend in the OFF state is *r/k*, and the expected number of generations a cell will spend in the ON state is *r/γ*.

When discussing the fraction of cells in the ON state, we will use *f* to refer to the average fraction of ON cells in the original population, and *F*(*t*) to refer to the fraction of cells in the ON state in cell colonies over time.

*f* can be written in terms of the state switching rates *k* and *γ* as

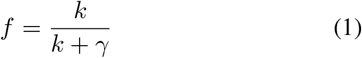

Individual cells are randomly chosen from the population, and they will be in the ON state with probability *f* and in the OFF state with probability 1 – *f*. Let *X*(*t*) and *Y*(*t*) denote the number of cells within the colony in the ON state and OFF state, respectively, at time *t*. The time evolution of integer-valued random processes *X*(*t*) and *Y*(*t*) is governed by the model illustrated in Table I.

**TABLE I.**
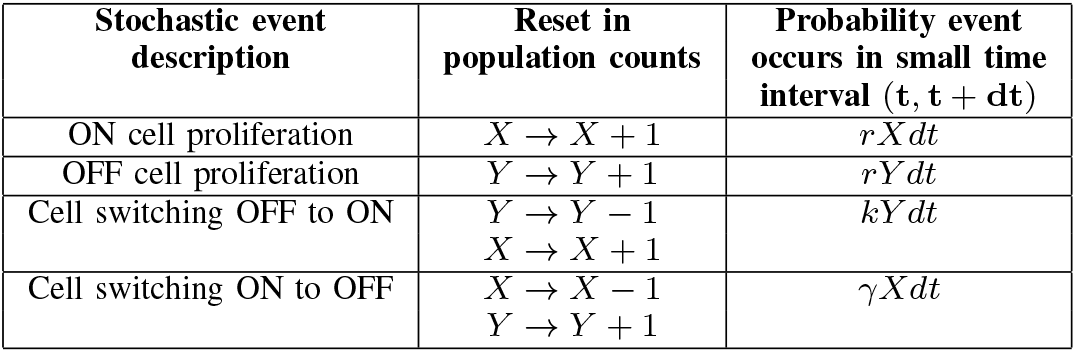
Stochastic Model of Cell Proliferation and Switching

The model consists of four events that occur probabilistically with rates given in the third column, and whenever the event occurs, the cell numbers increase/decrease by one as per the reset map in the second column. Based on this Markov process, we can derive the time evolution of the joint probability density function of *X*(*t*) and *Y*(*t*) that evolves according to the following Chemical Master Equation (CME)

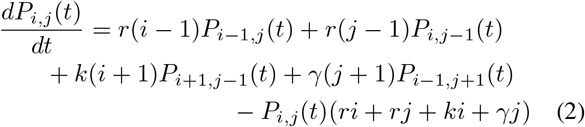

where *P_i,j_*(*t*) denotes the probability of observing *i* cells in the OFF state, and *j* cells in the ON state at time *t* [51], [52]. Previous research has shown how the time evolution of moments can be derived from the CME [53]-[56].

We are interested in the ratio

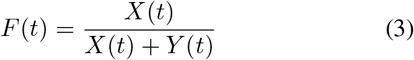

that represents the fraction of cells in the ON state at time *t*, and our goal is to quantify statistical fluctuations in this ratio across colonies. To do this, we derive the time evolution of the first two statistical moments of *X*(*t*) as well as those of the total cell count, which we will refer to as *T*(*t*) = *X*(*t*)+ *Y*(*t*). For the sake of simplicity, we will refer to the random variables *X*(*t*), *Y*(*t*), *T*(*t*), *F*(*t*), as simply *X, Y, T*, and *F* from now on.

We will use angle brackets to denote the expected value of a random variable. For example, 〈*X*〉 will denote the expected value of *X*, and 〈*X^m^*〉 Xwill denote the expected value of *X^m^*.

The following equation is a general formula for the moment dynamics ODE of this system [53].

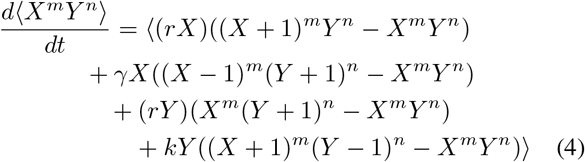

We can use this general formula to find differential equations for the first moments of *X* and *Y*, which are shown below.

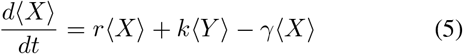

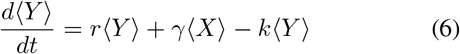

To define the initial conditions for this set of ODEs, we must consider the state of our experiment at time *t* = 0. At time *t* = 0, each colony is still only a single cell that has been sampled from the original population. If this individual cell is ON, then the fraction of cells in the ON state will be 1. If this cell is OFF, then the fraction of cells in the ON state will be 0. In other words, the fraction of cells in the ON state at time *t* = 0 follows a Bernoulli distribution, taking the value 1 with probability *p*(1) = *f*, and taking the value 0 with probability *p*(0) = 1 – *f*. So, at time *t* = 0, 〈*X*〉 = *f* and 〈*Y*〉 = 1 – *f*. With this information, we can solve this set of ODEs and find equations for 〈*X*〉 and 〈*Y*〉 as functions of time. These are shown below.

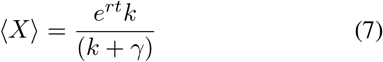

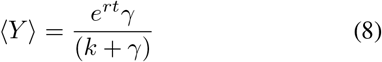

Adding equations (7) and (8) together yields an expression for 〈*T*〉, the total number of cells in both states.

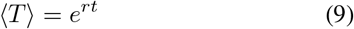

To compute the second moments of *X* and *Y*, we again use the differential equation technique described in equation (4). After simplifying, this yields:

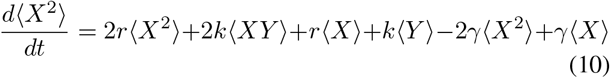

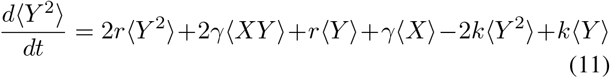

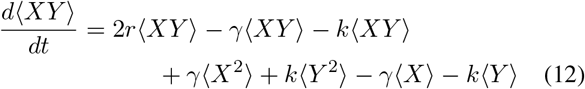

Again, we know the initial conditions for this set of ODEs: at time *t* = 0, 〈*X*^2^〉 = *f*, 〈*Y*^2^〉 = 1 – *f*, and 〈*XY*〉 = 0. Using this information, we can again solve these ODEs and find the second order moments as functions of time.

Equation (13), shown at the top of the next page because of its length, is the equation for 〈*X*^2^〉. We have omitted 〈*Y*^2^〉 and 〈*XY*〉 for now, as they are not needed for our fluctuation test analysis. To find the second moment of *T*, we will again use the differential equation method described in equation (4). However, this time, we will ignore state-switching dynamics and consider only cell population growth, since *T* is the total number of all cells in the population.

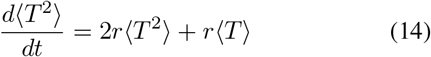

We know the initial condition at time *t* = 0 is 〈*T*^2^〉 = 1. After solving the ODE with this initial condition, the resulting equation for 〈*T*^2^〉 is shown below.

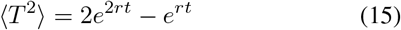

Based on equations (9) and (15), we note that the total number of cells *T* at a given timepoint follows a geometric distribution.

We have now derived equations for the first and second order moments of *X* and *T*. However, in order for this method to be experimentally and computationally feasible, what we are actually interested in is the first and second order moments of the fraction 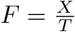, which is the fraction of cells in the ON state over time. If we can find the first and second order moments of this fraction, then we can use these moments to compute the *CV*^2^ of the fraction of ON cells across colonies as a function of time.

How can we compute the moments of *F*? Please consider the following:

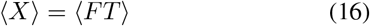

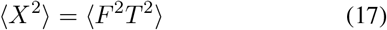

At this point, we will introduce a simplifying assumption that will allow us to derive an approximation of the moments 〈*F*〉 and 〈*F*^2^〉. We will assume that *F* and *T* are independent variables. In other words, the fraction of ON cells is independent of the total number of cells in the colony. In the next section of the paper, we will discuss how accurate this assumption is. Making this assumption, we can rewrite equations (16) and (17) as, respectively:

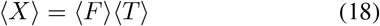

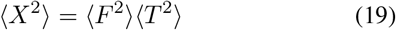

Then, simple algebra yields the first and second order moments of *F*:

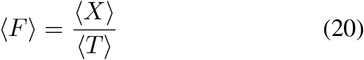

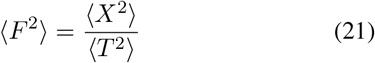

Please note that we have already derived 〈*X*〉, 〈*T*〉, 〈*X*^2^〉, and 〈*T*^2^〉 in equations (7), (9), (13), and (15) respectively. So, we can simply plug in those formulas to get the first and second order moments of *F* in terms of *k, γ*, and *t*. Equation (22) is the first moment of *F*, which is equal to *f*, the fraction of ON cells in the original population. Equation (23), shown on the following page because of its size, is the second moment of *F*.

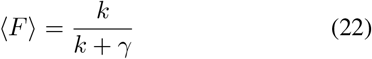

Finally, now that we have calculated the first and second order moments of the fraction of ON cells, we can calculate the coefficient of variation squared as follows:

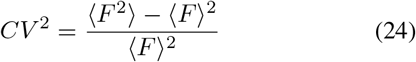

The full formula for *CV*^2^ is a bit long and unwieldy, so we will define two new parameters to be able to write it more simply. Let 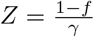, and let *τ* = *tr*, so that *τ* is the duration of the experiment normalized to generation time. Then we can write the formula for *CV*^2^ as follows.

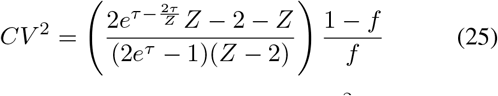

Equation 26 shows the limit of our *CV*^2^ formula as *t* approaches 0. To understand this, consider that at time *t* = 0, each colony is still only a single cell that has been sampled from the original population. So, the fraction of cells in the ON state will take the value 1 with probability *p*(1) = *f*, and will take the value 0 with probability *p*(0) = 1 – *f*. In other words, at time *t* = 0, the fraction of cells in the ON state follows a Bernoulli distribution, and the *CV*^2^ of this distribution is 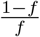.

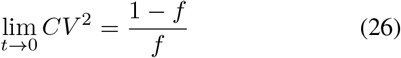

Conversely, as *t* goes to infinity, the fraction of ON cells in each of the colonies tends back towards the steady state value of *f* in the original population, and the *CV*^2^ of the fraction of ON cells in the colonies approaches 0.

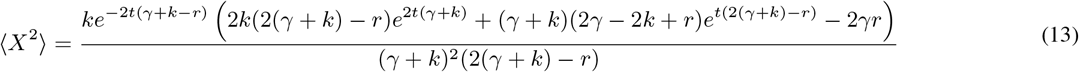

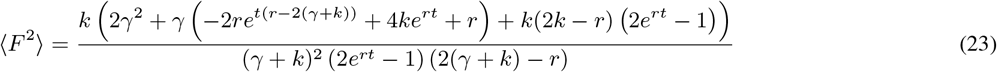

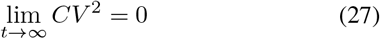

Importantly, at a fixed experimental endpoint *t*, cell populations with slower switching rates will tend to have a higher *CV*^2^ in the fraction of ON cells across colonies compared to cell populations with faster switching rates. The intuition behind this is that colonies from populations with faster switching rates will tend to converge back to the steady state fraction of ON cells more quickly, leading to less variation across colonies, compared those of a slower-switching population.

Figure 3 shows a comparison of our formula’s *CV*^2^ predictions and *CV*^2^ of stochastic simulations of the experiment. Now that we have a formula for the *CV*^2^ of the fraction of ON cells, we can measure the *CV*^2^ of the fraction of ON cells in real cell populations, and compare the formula to the real measurement to fit the state-switching parameters *k* and *γ*. Please note that, based on equation (1), if we experimentally measure the fraction of cells in the ON state in the original population, we can write *k* in terms of *γ* and this measured fraction.

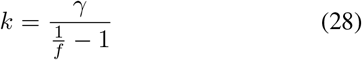

**Fig. 3.**
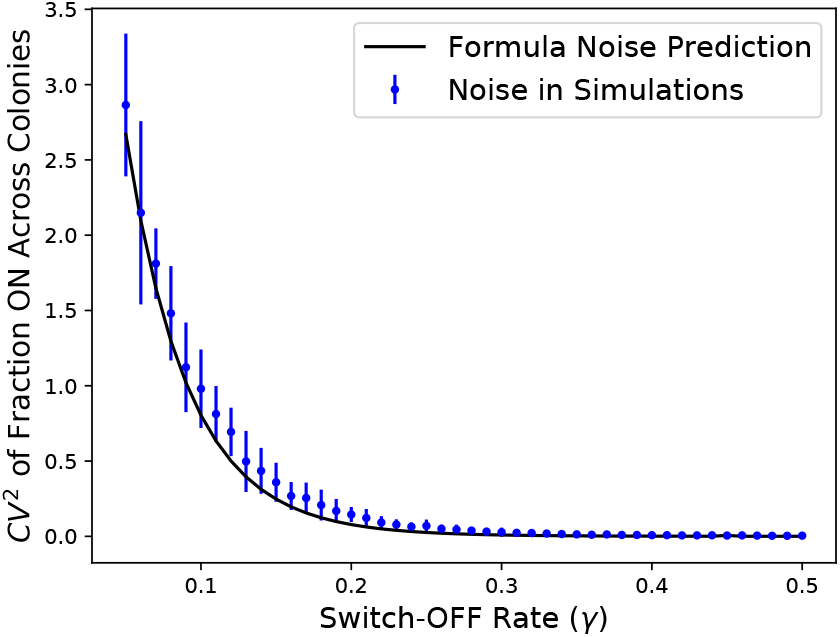
Comparison between formula noise prediction and noise in simulations. Simulated data was generated with the Gillespie algorithm method, further described in Section V. The fraction of ON cells in the original population (*f*) was assumed to be 0.10. We set the proliferation rate (*r*) to be 1, so that time is normalized to the generation time of the cells. The simulations were run for 12 generations (*t* = 12), and *CV*^2^ was calculated across 40 colonies. The simulations were run 50 times per parameter set to generate error bars, which show one standard deviation. Although our formula is not a perfect prediction of the true underlying *CV*^2^, it is a reasonably good approximation.

So, if we know the experimentally-measured *CV*^2^, the fraction of ON cells in the original population *f*, the duration of the experiment *t*, and have written *k* in terms of *γ*, then we are left with only one unknown variable, *γ*, that we can solve for with a simple error-minimization between the derived *CV*^2^ formula and the experimentally-measured *CV*^2^.

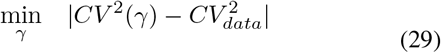

In practice, we use the Python SciPy library’s **optimize.fmin()** function to do this error minimization and solve for *γ*.

## IV. Independent Variable Assumption

As noted previously, in order to derive the formulas for the first and second order moments of *F*, we made a simplifying assumption: that *F* and *T* are independent variables (which would mean that *F*^2^ and *T*^2^ are also independent variables). To be precise, writing equation (16) as equation (18) is based on the assumption that *F* and *T* are independent, and writing equation (17) as equation (19) is based on the assumption that *F*^2^ and *T*^2^ are independent.

However, we tested this assumption by collecting data from stochastic simulations (discussed further in the next section), and found that it is not completely correct. For these simulations, we set the proliferation rate (*r*) to be 1, so that time is normalized to the generation time of the cells. The fraction of ON cells in the original population (*f*) was assumed to be 0.10. The switch-off rate (*γ*) was set at 1/8, so that the expected number of generations spent in the ON state is 8. The switch-on rate (*k*) was set according to equation (28), using these values of *f* and *γ*. The simulations were run for 7 generations (*t* = 7). We ran 10,000 simulations, and at the end of each one we recorded the total number of cells (*T*) and the fraction of cells in the ON state (*F*), as well as the squares of both (*F*^2^ and *T*^2^).

We observed a very slight Pearson correlation of −0.016 between *F* and *T*, which was not statistically significant (*p* = 0.1087). However, we observed a stronger Pearson correlation of −0.0439 between *F*^2^ and *T*^2^, which was statistically significant (*p* = 1.1078 × 10^-5^). This negative correlation is shown in Figure 4, a plot of the final fraction of cells in the ON state squared (*F*^2^) against the total number of cells squared (*T*^2^) for 10,000 simulated cell colonies. This observed statistical dependence between *F*^2^ and *T*^2^ means that there must also be some level of dependence between *F* and *T*, even if it was not detected to a statistically significant extent in our simulation analysis.

**Fig. 4.**
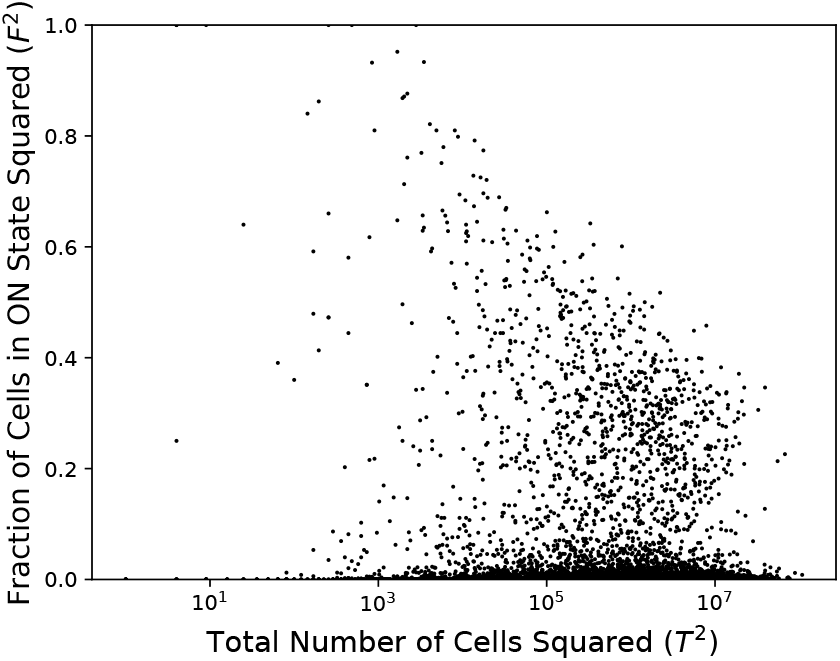
Fraction of cells in the ON state squared *F*^2^ plotted against the total number of cells squared *T*^2^ for 10,000 simulated cell colonies. For these simulations, we set the proliferation rate (*r*) to be 1, so that time is normalized to the generation time of the cells. The fraction of ON cells in the original population (*f*) was assumed to be 0.10. The switch-off rate (*γ*) was set at 1/8, so that the expected number of generations spent in the ON state is 8. The switch-on rate (*k*) was set according to equation (28), using these values of *f* and *γ*. The simulations were run for 7 generations (*t* = 7). We ran 10, 000 simulations, and at the end of each one we recorded the total number of cells squared (*T*^2^) and the fraction of cells in the ON state squared (*F*^2^). We observed a Pearson correlation of −0.0439 between *F*^2^ and *T*^2^, which was statistically significant (*p* = 1.1078 × 10^-5^).

So, although the derivation of our *CV*^2^ formula was based on a simplifying assumption of independence between *F* and *T*, it appears that there is in fact a very slight level of statistical dependence between them. Therefore, our derived formula for the *CV*^2^ should be thought of as an approximation, not as an exact analytical formula for the experimental *CV*^2^. Can this approximation of the *CV*^2^ still be useful in a biological context, even if it is not exact? In the following sections, we will benchmark our method on simulated data to test its efficacy.

## V. Simulating Cell Colonies

### A. Simulations with Gillespie Algorithm

In order to test if our fluctuation test method, based on the approximation of the *CV*^2^, is useful, we first generated simulated data by using the Gillespie algorithm [57] to simulate the stochastic model described previously in Table I.

Please note that for all of our simulations, we set the proliferation rate *r* = 1, so the average cell cycle time 1/*r* will also be 1 and time is normalized to the generation time of the cells. This will allow us to discuss results for 1/*γ* both as predicted time in the ON state and predicted number of generations in the ON state.

Figure 5 shows the evolution of the fraction ON, 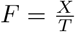, over time in the stochastic simulations (blue dotted curves), compared to the deterministic steady state fraction ON.

**Fig. 5.**
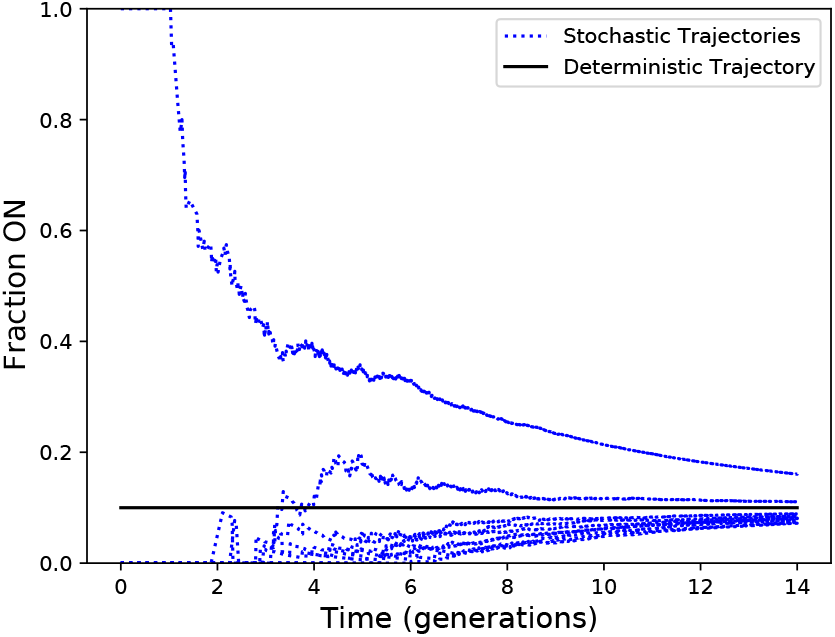
This plot shows the evolution of the fraction ON, 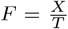, over time in the stochastic simulations (blue dotted curves), compared to the deterministic steady state fraction ON. The fraction of ON cells in the original population (*f*) was assumed to be 0.10. We set the proliferation rate (*r*) to be 1, so that time is normalized to the generation time of the cells. Most colonies begin with a single OFF cell, and the fraction ON increases up to 0.10 over time as the colony grows. A small number of colonies begin with a single ON cells, and the fraction ON decreases down to 0.10 over time as the colony grows.

To simulate a colony of cells, we will run this simulation starting with one cell that will be randomly chosen to either be in the *X* (ON) state with probability *f*, or the *Y* (OFF) state with probability 1 – *f*. Then we will run the simulation to some specified end point to simulate the colony of cells proliferating and switching between the ON and OFF states. To simulate the modified fluctuation test experiment, we will simulate many of these colonies so that we can measure the *CV*^2^ of the final fraction of ON cells in each colony.

### B. Non-Exponential Cell Cycle Time

In our simulations with the Gillespie algorithm, it is assumed that the timing between events follows an exponential distribution. However, in the biological context, cell cycle timing does not typically follow an exponential distribution. More realistic distributions for modeling cell cycle timing are the log-normal distribution and gamma distribution. So, in addition to using the Gillespie algorithm, we also developed an algorithm for simulating cell colonies that can account for other distributions of cell cycle timing besides the exponential distribution. More information about these simulations is available in the Appendix section of this paper, and the code for these simulations is available upon request from the authors.

## VI. Results

As discussed previously, the formula we derived for the *CV*^2^ of the fraction of ON cells across colonies over time is an inexact approximation. Is this approximation accurate enough to be useful to biologists in an experimental setting? To investigate this, we decided to benchmark our formula by generating simulated data and seeing how effective our method was at estimating the true, underlying rates of state-switching.

We considered two hypothetical cell populations: a fast-switching population and a slow-switching population. Both populations have an average of 10% ON cells in the original populations, so *f* = 0.10. However, cells in the fast-switching population spend an expected 4 generations in the ON state, while cells in the slow-switching population spend an expected 10 generations in the ON state, so *γ* = 1/4 for the fast-switching population and *γ* = 1/10 for the slow-switching population. We assumed that our simulated experiment would run for 12 generations (*t* = 12), and *CV*^2^ would be calculated across 40 colonies. We set the proliferation rate *r* = 1, so the average cell cycle time 1/*r* will also be 1 and time is normalized to the generation time of the cells, allowing us to discuss results for 1/*γ* both as predicted time in the ON state and predicted number of generations in the ON state.

We used the stochastic simulation methods described in the previous section to simulate the experimental workflow of the fluctuation test, generating many simulated datasets for both the fast-switching and slow-switching scenarios. We then applied our switch-rate estimation method to each of these “experimental” datasets, to see how close the predicted switching rates came to the actual switching rates. To make our results more intuitive, we have chosen to report 1/*γ*, which is the expected number of generations in the ON state, rather than *γ* itself.

The histograms in the following subsections show the predictions of 1/*γ* (expected number of generations in the ON state) for the fast-switching datasets in blue, and for the slow-switching datasets in red. There are also vertical lines at 4 and 10 on the horizontal axis, to mark the true, underlying values of 1/*γ* in the fast-switching and slow-switching scenarios, respectively.

### A. Exponential Distribution Cell Cycle Time

Figure 6 shows the results of the benchmarking analysis, for cell colonies simulated with the Gillespie algorithm, assuming exponentially-distributed cell cycle time. As you can see, our method’s predictions for the fast-switching scenario cluster around the true 1/*γ* value of 4 (meaning that cells spend an average of 4 generations in the ON state), while the predictions for the slow-switching scenario cluster around the true 1/*γ* value of 10 (meaning that cells spend an average of 10 generations in the ON state). Table II shows relevant statistics for the predictions. The mean predicted 1/*γ* value for the fast-switching scenario was 4.67, while the mean predicted 1/*γ* value for the slow-switching scenario was 11.01.

**Fig. 6.**
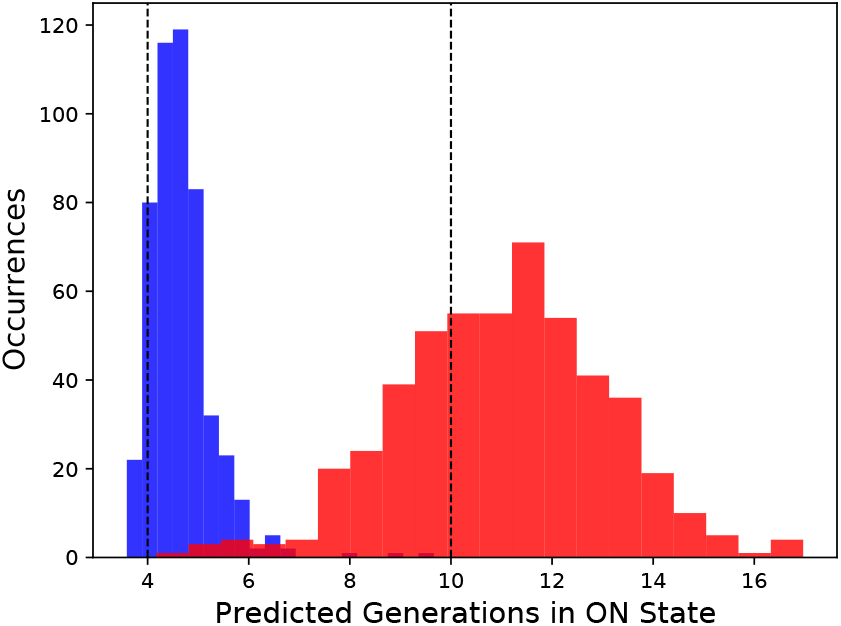
Benchmarking with exponentially-distributed cell cycle time, using data generated with the Gillespie algorithm. The fraction of ON cells in the original population (*f*) was assumed to be 0.10. We consider a fast-switching scenario, in which cells spend an average of 4 generations in the ON state (1/*γ* = 4), and a slow-switching scenario, in which cells spend an average of 10 generations in the ON state (1/*γ* = 10). We set the proliferation rate (*r*) to be 1, so that time is normalized to the generation time of the cells. The simulations were run for 12 generations (*t* = 12), and *CV*^2^ was calculated across 40 colonies. Our method’s predictions for the fast-switching scenario (shown in blue) cluster around the true 1/*γ* value of 4 (shown with a dotted line), while the predictions for the slow-switching scenario (shown in red) cluster around the true 1/*γ* value of 10 (shown with a dotted line).

**TABLE II.** Benchmarking Results – Exponential Distribution of Cell Cycle Time

| Scenario | True Generations ON | Predicted Generations ON | Prediction Std. Dev. |
| --- | --- | --- | --- |
| Fast-Switching | 4 | 4.67 | 0.63 |
| Slow-Switching | 10 | 11.01 | 2.03 |

For both the fast-switching and slow-switching scenarios, our formula had a tendency to slightly overestimate the number of generations in the ON state. So, its performance was not perfect. However, we believe that it was still close enough to potentially be useful to biologists an approximation.

### B. Log-Normal Distribution Cell Cycle Time

Figure 7 shows the results of the benchmarking analysis, assuming a log-normal distribution of cell cycle time. Again, we observe that our method’s predictions for the fast-switching scenario cluster around the true 1/*γ* value of 4, while the predictions for the slow-switching scenario cluster around the true 1/*γ* value of 10. The mean predicted 1/*γ* value for the fast-switching scenario was 4.77, and the mean predicted 1/*γ* value for the slow-switching scenario was 10.97 (listed in Table III). Just as with the results in the previous sub-section, our formula again had a tendency to slightly overestimate the number of generations in the ON state in both the fast-switching and slow-switching scenarios. Again, we note that although our formula is not perfect, it may still be accurate enough to be useful to a biologist as an approximation.

**Fig. 7.**
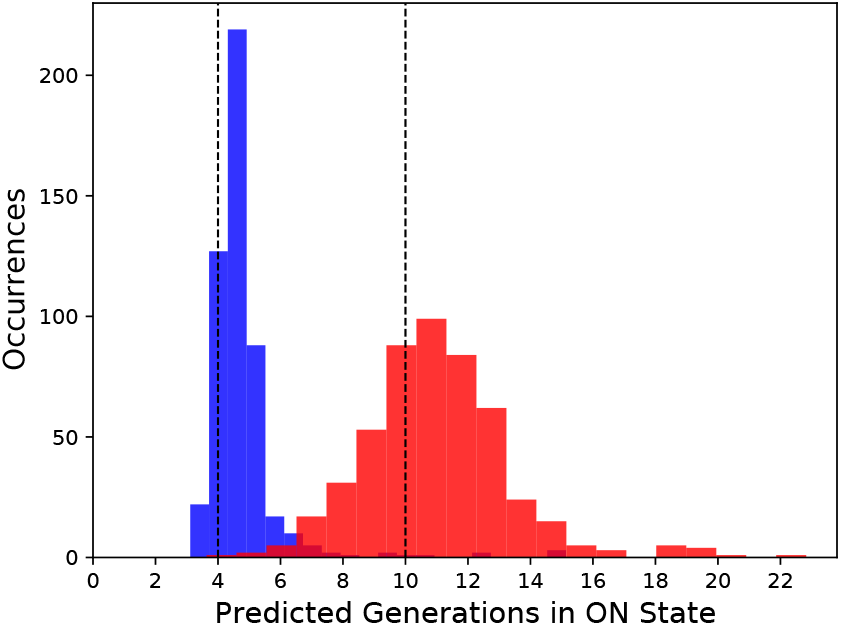
Benchmarking with log-normal-distributed cell cycle time. The fraction of ON cells in the original population (*f*) was assumed to be 0.10. We consider a fast-switching scenario, in which cells spend an average of 4 generations in the ON state (1/*γ* = 4), and a slow-switching scenario, in which cells spend an average of 10 generations in the ON state (1/*γ* = 10). We set the proliferation rate (*r*) to be 1, so that time is normalized to the generation time of the cells. The simulations were run for 12 generations (*t* = 12), and *CV*^2^ was calculated across 40 colonies. Our method’s predictions for the fast-switching scenario (shown in blue) cluster around the true 1/*γ* value of 4 (shown with a dotted line), while the predictions for the slow-switching scenario (shown in red) cluster around the true 1/*γ* value of 10 (shown with a dotted line).

**TABLE III.** Benchmarking Results – Log-Normal Distribution of Cell Cycle Time

| Scenario | True Generations ON | Predicted Generations ON | Prediction Std. Dev. |
| --- | --- | --- | --- |
| Fast-Switching | 4 | 4.77 | 1.23 |
| Slow-Switching | 10 | 10.97 | 2.33 |

### C. Gamma Distribution Cell Cycle Time

Figure 8 shows the results of the benchmarking analysis, assuming a gamma distribution of cell cycle time. The mean predicted 1/*γ* value for the fast-switching scenario was 4.67, and the mean predicted 1/*γ* value for the slow-switching scenario was 10.87 (listed in Table IV). Just as with the exponential distribution and log-normal distribution simulations, we again observe a tendency of our formula to slightly overestimate the true number of generations in the ON state. However, yet again the formula estimates of the number of generations ON are close enough to the true underlying values to potentially be useful in a biological context, even though they are not exact.

**Fig. 8.**
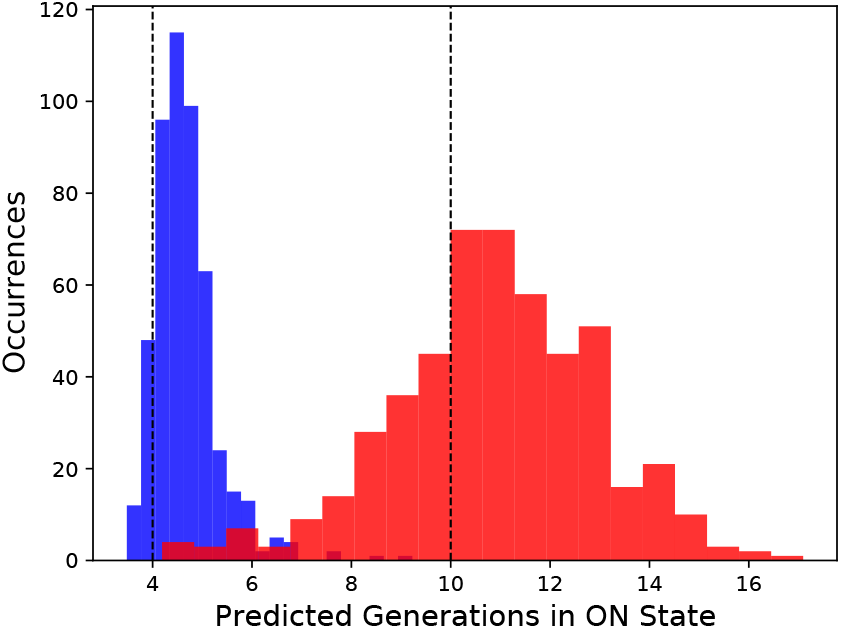
Benchmarking with gamma-distributed cell cycle time. The fraction of ON cells in the original population (*f*) was assumed to be 0.10. We consider a fast-switching scenario, in which cells spend an average of 4 generations in the ON state (1/*γ* = 4), and a slow-switching scenario, in which cells spend an average of 10 generations in the ON state (1/*γ* = 10). We set the proliferation rate (*r*) to be 1, so that time is normalized to the generation time of the cells. The simulations were run for 12 generations (*t* = 12), and *CV*^2^ was calculated across 40 colonies. Our method’s predictions for the fast-switching scenario (shown in blue) cluster around the true 1/*γ* value of 4 (shown with a dotted line), while the predictions for the slow-switching scenario (shown in red) cluster around the true 1/*γ* value of 10 (shown with a dotted line).

**TABLE IV.** Benchmarking Results – Gamma Distribution of Cell Cycle Time

| Scenario | True Generations ON | Predicted Generations ON | Prediction Std. Dev. |
| --- | --- | --- | --- |
| Fast-Switching | 4 | 4.67 | 0.65 |
| Slow-Switching | 10 | 10.87 | 2.06 |

### D. Conclusion

Overall, we conclude that this method of switch-rate estimation is efficacious enough to be useful in certain contexts. Our formula is imperfect and has a tendency to slightly overestimate the number of generations spent in the ON state. However, it is close enough to potentially be useful to biologists as an approximation. Also, our formula has the advantage of being robust to different distributions of cell cycle time. This is a non-trivial result, since the moment analysis techniques used to derive our formula typically assume an exponential distribution of time between events in the simulation, and is an important advantage in real biological contexts, in which the underlying distribution of cell cycle timing may not be known.

## VII. Discussion

In this paper, we described a modified fluctuation test that can be done to estimate the rates of transient state-switching in cell populations. We analyzed a stochastic model of the experiment to derive a moment-based formula for the *CV*^2^ of the fraction of cells in the ON state over time. One step in this derivation involved making a simplifying assumption, that the fraction of cells in the ON state *F* and the total number of cells *T* are independent, uncorrelated variables. This assumption is not completely correct. In fact, there is a very slight negative correlation between these variables. So, our formula is not exact, but rather an approximation. Despite this imperfection, we wondered if this approximation could still be useful. We benchmarked it on simulated cell populations, using varied assumptions for the distribution of cell cycle time. We found that our method was able to differentiate between fast-switching and slow-switching simulated cell populations to a reasonable degree, suggesting that it may be a useful tool for biologists despite its imperfection.

That the method works well for simulations with gamma-distributed and log-normal-distributed cell cycle times is a non-trivial result, since the moment analysis techniques used to derive the formula assumed exponentially-distributed time between events. Being robust to many potential distributions of cell cycle time is an important advantage, since in a real biological setting the exact underlying distribution of the cell cycle time may not be known.

Future work on this topic will include applying our method to real biological datasets in collaboration with experimental biologists working on topics such as drug resistance in cancer. Also, we will further explore the theory of this method by relaxing some of the assumptions in our problem formulation. For example, in this paper we assumed that cells in both states grow at the same rate, so it will be interesting to explore the theoretical implications of differences in growth rates between the states. Furthermore, we hope to expand this model beyond two states to study systems with three or more states.

## Appendix

For readers who are not interested in computer programming or familiar the Python language, this section can be skipped. Just note that we used a simulation of cell colonies in which cells switch back and forth between two states, and divide with cell cycle times that can follow log-normal or gamma distributions. However, for readers interested in computer programming and familiar with Python, we will now describe our simulation code. please note that this code is available upon request from the authors. The Python code is listed below:

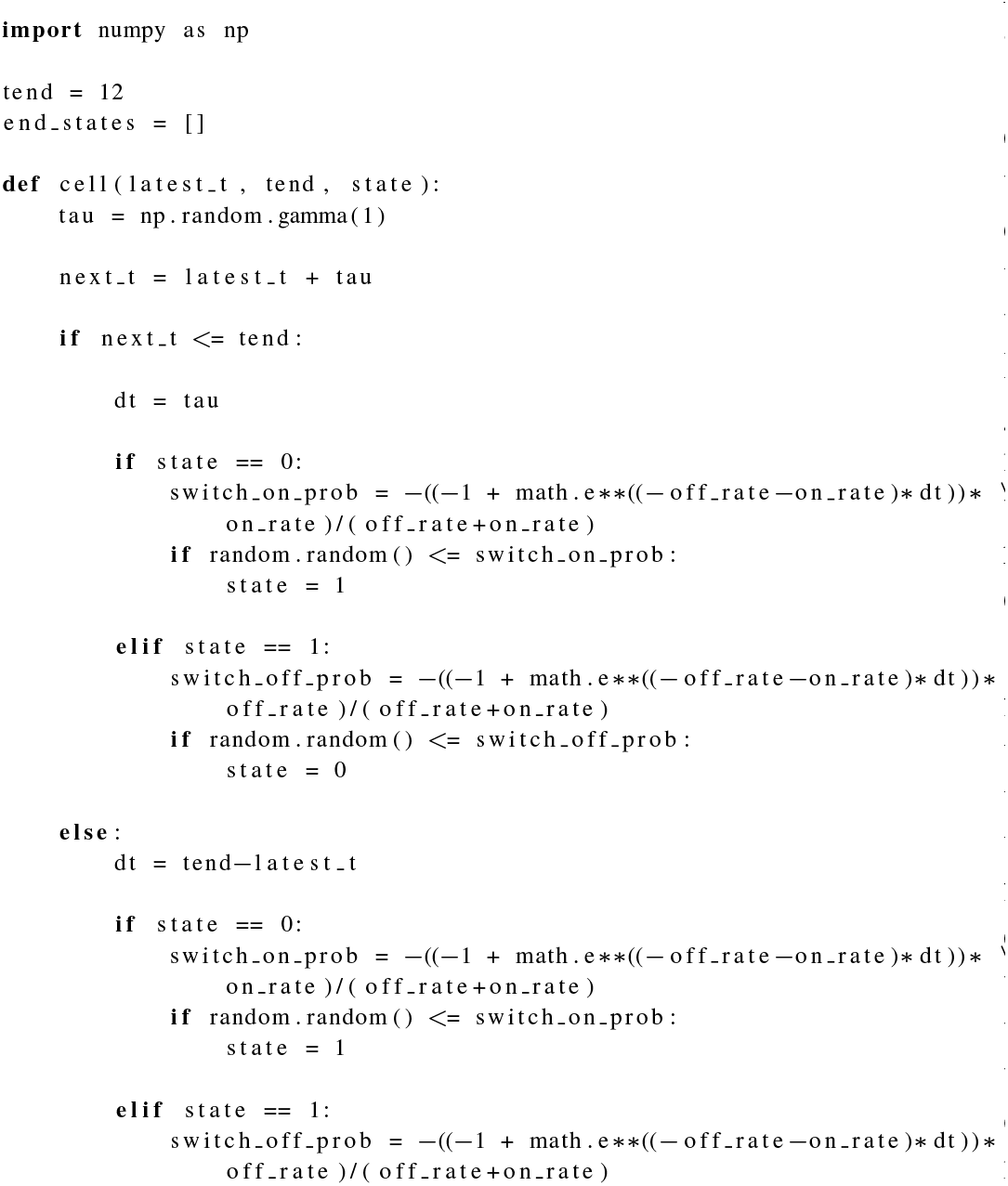
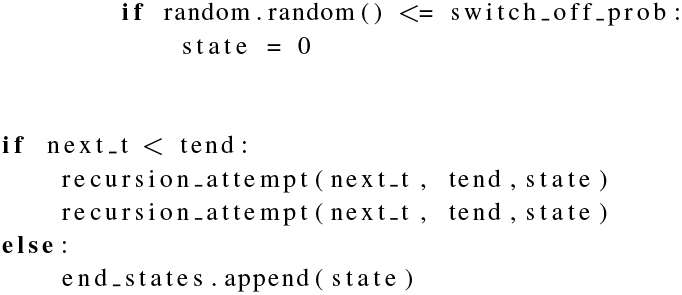

This simulation algorithm is based on a recursive function **cell()** which represents a single cell. The function takes three arguments: **latest_t** is the timepoint at which the cell is instantiated, **tend** is the global variable that defines the end time of the simulated experiment, and **state** is the state of the cell, where 0 represents the OFF state and 1 represents the ON state.

After a cell is instantiated, we then determine how long it will take for the cell to divide by taking a random draw from some probability distribution, and save this in the local variable **tau**. Importantly, we can choose to draw this number from any distribution, and are not limited to the assumption of exponential cell cycle time. In the example code, we are drawing from a gamma distribution with a mean of 1, using the NumPy library, which we have imported at the top of the code.

Once we have chosen the cell cycle time of this cell, we must check to see if it switches states before dividing. To do this, we have derived formulas for the probability of being ON after some time *t* if a cell starts in the OFF state, and vice versa. Please note that a cell could theoretically switch back many times before dividing, so in order to calculate these probability formulas it was necessary to use the moment analysis techniques described in Section III of this paper.

Next, we check to see if the time of division of this cell exceeds the end time of the experiment **tend**. If it doesn’t, then we recursively call the function to instantiate two new cells, representing the daughter cells of the original cell, with their birth timepoints being the division timepoint of the original cell, and inheriting the state of the original cell. However, if the division timepoint of the original cell is greater than the **tend** endpoint of the experiment, then we do not call the function again, and instead append the state of the cell to the global list **end_states**, which will eventually record the state of every cell in the colony at the end of the experiment.

So, we can simulate a colony of cells by calling the **cell()** function with a start-time of 0, an end time of whatever we set as our **tend**, and a randomly-selected initial state, which will be 1 (ON) with probability *f*, and 0 (OFF) with probability 1 – *f*. After all recursive iterations of the functions are finished running, it will leave us with our **end_states** full of the end states of each cell in the colony at the end of the experiment. In order to do a full simulation of the modified fluctuation test experiment, we can simply run this colony simulation many times, and record the fraction of cells in the ON state each time, which will allow us to measure the *CV*^2^ in these fractions across colonies.

## ACKNOWLEDGMENTS

This work is supported by grants from the Army Research Office (W911NF1910243) and the National Science Foundation (ECCS-1711548).

